# Astrocyte sensitivity to glymphatic shear stress is amplified by albumin and mediated by the interaction of sphingosine 1 phosphate with Piezo1

**DOI:** 10.1101/2023.11.06.565884

**Authors:** David Ballesteros-Gomez, Sean McCutcheon, Greta L. Yang, Antonio Cibelli, Ashley Bispo, Michael Krawchuk, Giselle Piedra, David C. Spray

## Abstract

Astrocyte endfeet enwrap brain vasculature, forming a boundary for perivascular glymphatic flow of fluid and solutes along and across the astrocyte endfeet into the brain parenchyma. To determine whether astrocytes may sense and respond to the shear forces generated by glymphatic flow, we examined intracellular calcium (Ca^2+^) changes evoked in astrocytes to brief fluid flow applied in calibrated microfluidic chambers. Shear stresses < 20 dyn/cm^2^ failed to evoke Ca^2+^ responses in the absence of albumin, but cells responded to shear stress below 1 dyn/cm^2^ when as little as 5 μM albumin was present in flow medium. A role for extracellular matrix in mechanotransduction was indicated by reduced sensitivity after degradation of heparan sulfate proteoglycan. Sphingosine-1-phosphate (S1P) amplified shear responses in the absence of albumin, whereas mechanosensitivity was attenuated by the S1P receptor blocker fingolimod. Piezo1 participated in the transduction as revealed by blockade by the spider toxin GsMTX and amplification by the chemical modulator Yoda1, even in absence of albumin or S1P. Our findings that astrocytes are exquisitely sensitive to shear stress and that sensitivity is greatly amplified by albumin concentrations encountered in normal and pathological CSF predict that perivascular astrocytes are responsive to glymphatic shear stress and that responsiveness is augmented by elevated CSF protein. S1P receptor signaling thus establishes a setpoint for Piezo1 activation that is finely tuned to coincide with albumin level in CSF and to the low shear forces resulting from glymphatic flow.

**Graphical abstract:** 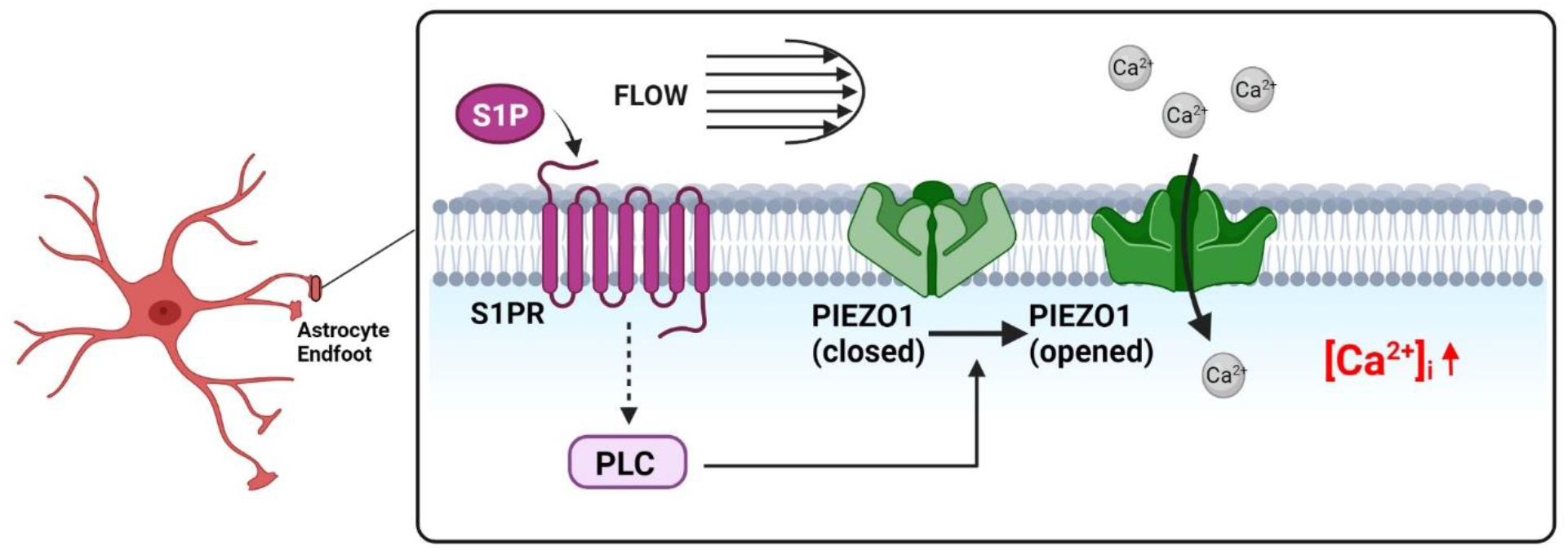

Astrocyte endfoot responds to glymphatic shear stress when albumin is present. Mechanism involves sphingosine-1-phosphate (S1P) binding to its receptor (S1PR), activating phospholipase C (PLC) and thereby sensitizing the response of Piezo1 to flow. Ca^2+^ influx triggers Ca^2+^ release from intracellular stores and further downstream signaling, thereby modulating parenchymal perfusion. Illustration created using BioRender.com

## Introduction

Neural cells are separated from the brain circulation by a multilayered barrier of vascular endothelial cells and smooth muscle or pericytes, a perivascular space that appears as thin basement membrane in electron micrographs, and astrocyte endfoot processes that enwrap the vessels (Iadecola, 2017). Astrocyte endfeet release vasoactive factors that control vessel tone and thus dictate local blood flow, a process termed functional hyperemia (MacVicar and Newman, 2015, Anderson and Nedergaard, 2003). They are enriched in channels and transport mechanisms that take up water and solutes and thereby drive perfusion of the brain parenchyma (Simard and Nedergaard, 2004)(MacAulay, 2021).

Cerebrospinal fluid (CSF) flows from subarachnoid sinuses through perivascular spaces between vascular cells and astrocytes, forming the so-called glymphatic circulation (Iliff et al., 2012, Mestre et al., 2020). The topology, volume, and content of perivascular space through which the glymphatic circulation flows have been subjects of considerable experimental research and hydraulic modeling studies (e.g., (Min Rivas et al., 2020)). Whereas electron microscopy has revealed a basement membrane between endothelial cells and astrocyte endfeet that is generally quite thin (< 0.5 μm) and filled with dense matrix material (Reed et al., 2019), this narrow space reflects collapse as a consequence of fixation and its occlusion is at odds with functional studies showing dilated spaces for diffusion and rapid trajectories of perivascular nanoparticles (Min Rivas et al., 2020). Forces on the astrocyte endfeet that are generated by the pulsations of vasculature and resultant flow within the perivascular space include strain imposed by vascular constriction and dilation, and interstitial flow between the endfeet from basement membrane to parenchyma (see (Li et al., 2010)). In addition, because the astrocyte endfoot creates a boundary for glymphatic flow, shear stress is expected to result from the flow trajectories that have been visualized by perivascular nanoparticle movements (Plog et al., 2018).

Previous studies have measured astrocyte Ca^2+^ responses to vascular pressure gradients focally applied in acute brain slices (Kim and Filosa, 2012, Kim et al., 2016) and to brief concussive forces applied to astrocyte cell cultures (Ravin et al., 2019, Maneshi et al., 2015), concluding that astrocytes are sensitive to mechanical stimuli but require relatively large forces in order to respond. However, astrocyte sensitivity to low magnitude shear stresses corresponding to normal glymphatic flow have not been quantitatively examined. Moreover, previous flow studies on astrocytes have used “artificial CSF” and other simple saline solutions that lack proteins such as albumin that are normally present at low concentrations in normal CSF. We have examined the sensitivity of astrocytes to low levels of mechanical stimuli using primary cultures of mouse astrocytes, in which we quantified intracellular Ca^2+^ responses to a range of calibrated shear forces in flow chambers with solutions containing CSF levels of protein. These experiments revealed that astrocytes are quite sensitive to shear stress when evaluated in the presence of physiological albumin levels. Notably, the threshold albumin concentration is very nearly the same as normal CNS protein levels, and the threshold force detected by astrocytes under these conditions is consistent with flow rates estimated from perivascular movements of nanoparticles in vivo. We conclude that astrocyte endfeet directly sense and respond to shear stress generated by glymphatic flow, and that sensitivity to these low flow rates is greatly enhanced under conditions of heightened albumin levels as occur following breach of the blood brain barrier, in response to infection, or consequent to autoimmune disorders.

## Methods

### Primary Astrocyte Isolation and cell culture

Experiments were performed entirely on primary cultures of neonatal murine astrocytes, isolated as described (Cibelli et al., 2021). In brief, perinatal (Postnatal days 1-3) C57BL/6J (Jackson Labs #000664) or GCaMP6 mice (crossbreeds of homozygous Gfap-Cre (B6.Cg-Tg(Gfap-cre) 73.12 Mvs/J), Jackson Labs #012886) and GCaMP6f ^(sB6J.Cg-Gt(ROSA)26Sortm95.1(CAG-GCaMP6f)Hze/MwarJ,Jackson^ Labs #028865 strains) were euthanized via decapitation and cortical tissue was isolated by excision and removal of meninges in ice cold phosphate buffered saline (PBS) (Corning #21-030-CV). Isolated cortices ^f^were digested in 0.05% trypsin (Gibco #25200-056) for 10 min. Digested tissue was replated in tissue culture treated flasks (Falcon 353104) in astrocyte media: DMEM (Corning #10-014-CV) supplemented with 10% fetal bovine serum (FBS) (Thermo #10437-028) and 1% penicillin-streptomycin (PS) (Gibco #15070-063). At 14 days, flasks were shaken at 180 rpm for 12-24h in an incubated shaker (Ecotron HT) to remove residual tissue and microglia. Astrocytes were cultured up to passage 3 for use in experiments.

### Antibodies

The following primary antibodies were used: mouse monoclonal anti-GFAP (1:500; Sigma cat # SAB5201104), mouse monoclonal anti-HSPG (1:200; US Biological cat # H1890), rabbit polyclonal anti-Laminin (1:50; Santa Cruzcat # sc20142) and mouse monoclonal anti-CSPG (1:200; Sigma cat # C8035). Phalloidin Alexa Fluor 594–conjugated (1:1000; Thermo Fischer cat # A12381) was used to stain F-Actin.

The secondary antibodies used for immunofluorescence analysis were donkey anti-mouse Alexa Fluor 488–conjugated (1:1000; Invitrogen cat # A21202), donkey anti-mouse Alexa Fluor 594– conjugated (1:1000; Invitrogen cat # A-21203) and donkey anti-rabbit Alexa Fluor 488-conjugated (1:1000; Invitrogen cat # A-21206).

### Immunofluorescence

To assess the relative purity of primary astrocytes culture, cells were fixed with 4% paraformaldehyde for 15 minutes, washed 3 times in PBS, and permeabilized with 0.3% Triton X-100. After blocking using 2% BSA for 30 minutes, cells were incubated for 2 hours with primary antibodies at room temperature and washed with PBS BSA. Cells were finally incubated with secondary antibodies and mounted with ProLong (Invitrogen, cat# P36965), and DAPI for nuclear staining.

To evaluate the removal of cell surface proteoglycans by treatment with heparinase III (0.25U/mL for 2 hours) or chondroitinase ABC (0.2 U/mL for 2 hours), both controls and enzyme exposed cells were immunostained as previously described (MM Thi et al., 2004). Briefly, cells were chilled on ice and washed 3 times with ice cold PBS. Cells were then blocked with 1% BSA and incubated on ice with primary antibodies for 2 h, followed by ice-cold PBS BSA rinses and fixation with 4% paraformaldehyde. Cells were finally incubated with secondary antibodies and mounted as described above.

### Confocal microscopy and image analysis

Confocal images were obtained with an automated inverted Leica DMi8 SP8 microscope using a 63X NA = 1.4 Oil PL APO, WD = 0.14 mm objective. All confocal images were collected using 594 and 488 nm laser lines for excitation and a pinhole diameter of 1 Airy unit. Images were taken serially from top to bottom of each cell field with a raster size of 1024 × 1024 in the x–y planes and a z-step of 0.2-0.5 μm between optical slices (typically 15-25 optical sections per stack relative to different protein staining). Leica Application Suite X (Leica Microsystems CMS GmbH) and Fiji-ImageJ software were used to construct and process three-dimensional (3D) surface plot images and projections from z-stack.

The purity of primary astrocytes culture was quantitatively analyzed by identifying the nuclei overlapped with GFAP. The analysis was conducted on 3 to 5 different ROI from each of 3 independent experiments. All the cells in each field were counted using Fiji-ImageJ software and analyzed using GraphPad Prism 9.

To quantify the extent of HSPG and CSPG removal by proteases, immunostained-positive cells were identified and at least four different square regions of interest (ROI; fixed squared 10 μm^2^) per cell were selected. The average protein fluorescence intensity profile for each ROI was plotted and measured by using Fiji-ImageJ software and analyzed using GraphPad Prism 9 software for statistical analysis.

### Parallel plate flow experiments

Primary astrocytes up to passage 3 were plated in Ibidi-coated parallel plate flow chambers (micro-SlideVI^0.4^, ibidi #80606) 24-48 hours prior to experiments and maintained in astrocyte media (see above). One hour prior to testing, cells were loaded with 10 µM Fura-2 AM (Thermo #F1221) in astrocyte culture media. Post-incubation with Fura-2 AM, flow chambers were connected to syringe pumps (New Era model NE 300) via 20 or 60 mL syringes and chambers were rinsed for 1-3 min at 0.3 mL/min with flow media. Flow solutions were prepared by dilution of bovine serum albumin (30% BSA stock; VWR #VK719) to 0.3% (45 μM) or other desired concentration at the time of the experiment in extracellular solution containing 120 mM NaCl, 4 mM KCl, 2 mM CaCl_2_, 2 mM MgCl_2_, 10 mM HEPES, 10 mM D-glucose, pH 7.4 (https://cshprotocols.cshlp.org/content/2016/8/pdb.rec093229.short)

Applied shear stresses (ranging from <0.1 to >20 dyn/cm^2^) were calculated according to formulae in Application Note 11: Shear Stress and Shear Rates for ibidi μ Slides (ibidi GmbH, Version 4.1, 29 March 2016) and validated in initial experiments by weighing triplicate flow samples encompassing the range of flow rates used. For experiments involving change of solutions, flow was switched via stopcock valve between two syringe pumps, connected to the microfluidic chamber by low dead volume tubing. Cells were imaged by Nikon microscope (TE 2000, 20x 0.65 NA objective) with MetaFluor software (Molecular Devices) using Hamamatsu camera and Sutter filter changer connected to the microscope via a fiber optic cable. Intracellular Ca^2+^ was measured at 2 Hz by imaging regions of interest corresponding to individual astrocytes (diagramed in Fig 1A) with 380/340 nm dual excitation (Sutter filter changer) and Fura-2 filter cube. Ratio values were converted to intracellular Ca^2+^ concentrations as in our previous studies (Thi et al., 2013b). In some experiments, Ca^2+^ changes were measured as change in fluorescence intensity using 488 nm excitation and FITC filter cube in response to flow in astrocytes cultured from GFAPCre::GCaMP6f mice (see Fig S1).

**Figure 1:**
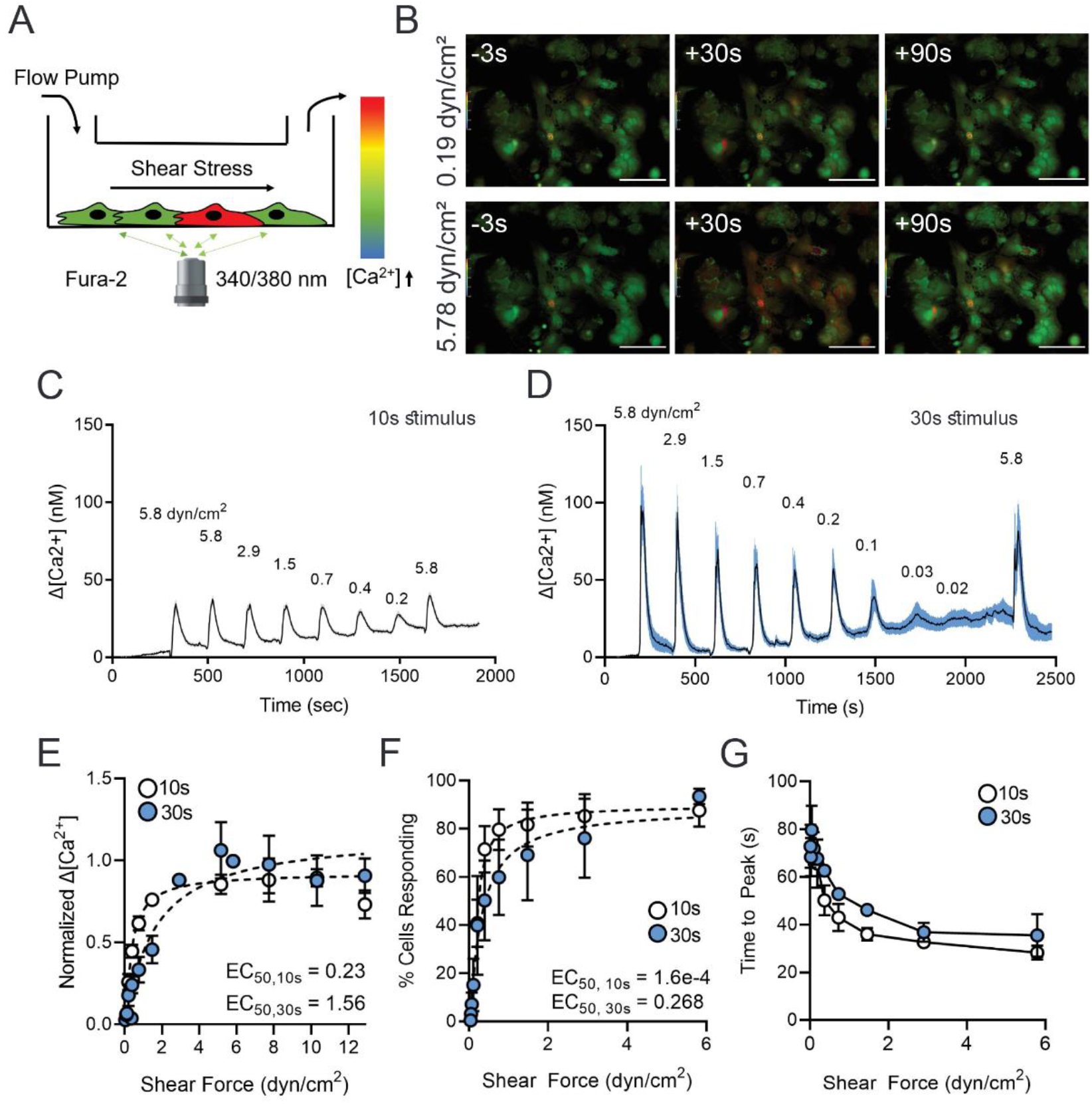
Astrocytes are highly sensitive to shear stress. (A) Diagram of dual-wavelength Fura-2 imaging of astrocytes in parallel flow chambers. (B) Representative ratiometric fluorescent micrographs of Ca^2+^ response time course in astrocytes at moderate (0.19 dyn/cm^2^) and high (5.78 dyn/cm^2^) shear forces. Scale bars 100 µm. (C) Representative change in [Ca^2+^]_i_ for a series of shear stresses using 10s stimuli and (D) using 30s stimuli. (E) Normalized change in Ca^2+^ concentration amplitude across shear rates with 10s and 30s stimuli. (F) Percentage of cells responding to a range of shear rates for 10s and 30s stimuli. (G) Time from stimulus onset to peak Ca^2+^ concentration across a range of shear stresses for 10s and 30s stimuli. All data shown as mean ± SEM. n ≥ 3 experiments for all treatments.

In addition to a few experiments examining high flow rates and in which we compared responses to varied stimulus duration at constant flow (Fig S2), two protocols were routinely used to quantify sensitivity to flow. In both protocols, stimuli were generally separated by 5 min intervals. Protocol 1 consisted of 10s or 30s exposures to moderate flow (4.5-5 ml/min), followed by progressive two-fold flow decreases to 0.1 ml/min or less, followed by another 4.5-5 ml/min stimulus (corresponding to a range from 5.8-6.6 to <0.2 dyn/cm^2^ as labeled in raw data figures and analysis) at 3 min intervals. As illustrated in Fig S3, similar responses were observed regardless of whether flow rates were decreased from high to low or increased from low to high. Protocol 2 consisted of paired moderate flow stimuli presented in a sequence of two 10 s or alternating 10s and 30s durations, wash in/out BSA or drug at 0.3 ml/min for 2 min, paired stimulation, wash in/out BSA or drug at 0.3 ml/min for 2 min, then paired stimulation (See Fig 2D). BSA experiments were performed with BSA concentrations of 0, 0.05, 0.1, 0.3, and 0.6% (0-90 μM) using both Protocols 1 and 2.

**Figure 2:**
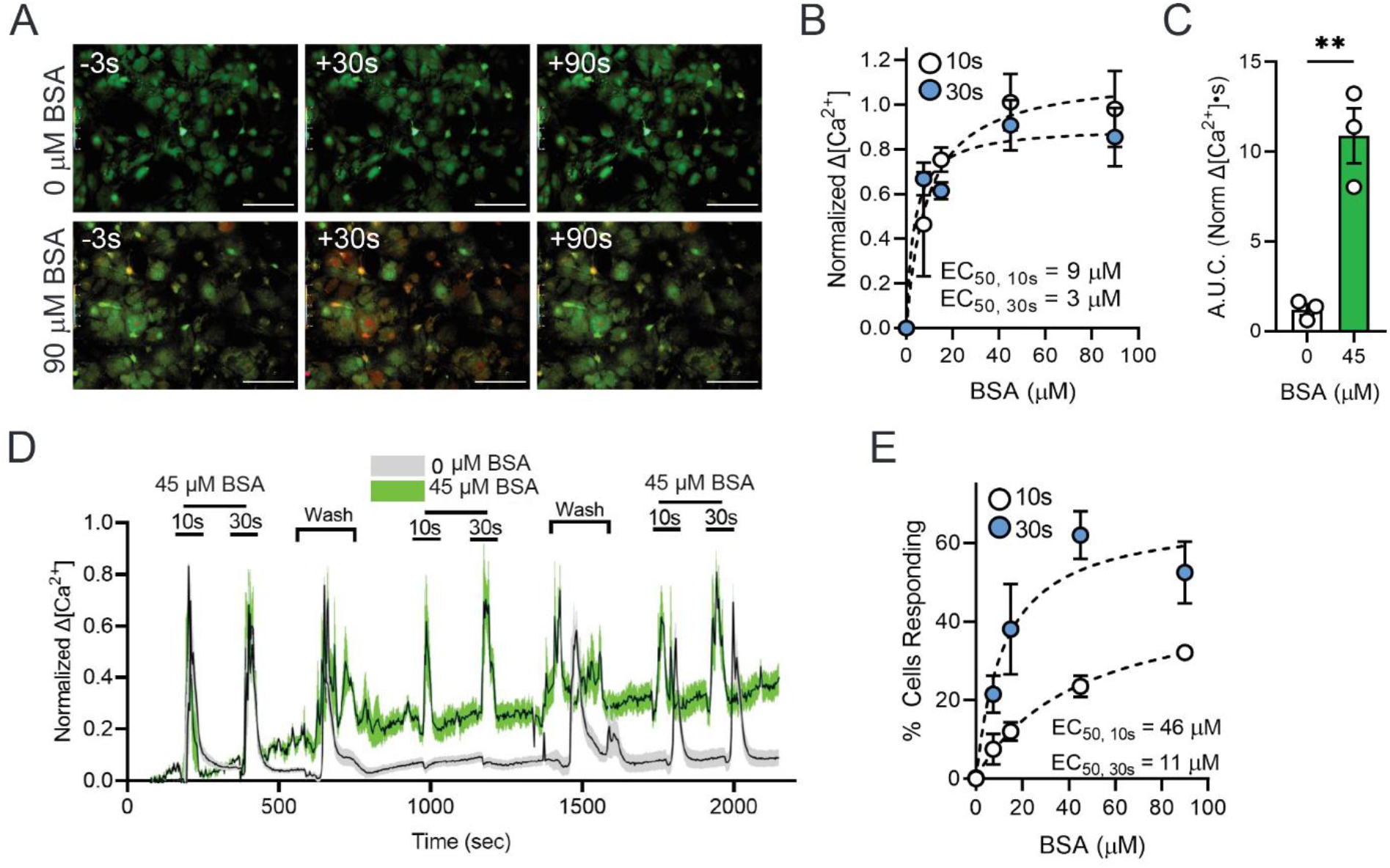
Astrocyte response to shear stress is dependent on albumin concentration. (A) Representative ratiometric fluorescence micrographs of Ca^2+^ response time course in astrocytes at 0and 90 M BSA using 2 mL/min flow rate (2.57 dyn/cm^2^). Scale bars 100 µm. (B) Normalized change in Ca^2+^ concentration amplitude across shear rates with 10s and 30s stimuli. (C) Area under normalized Ca^2+^ response curve (AUC) for 30s stimuli in 0 and 45 BSA. Unpaired Student’s t test: **p < 0.01. (D) Representative traces of normalized change in Ca^2+^ concentration in 0 and 45 BSA. (E) Percentage of cells responding to a range of shear rates for 10s and 30s stimuli. All data shown as mean ± SEM. n ≥ 3 for all experiments.

Fluorescence ratios were generally quantified in >20 individual ROIs in each experiment, and values presented in histograms represent means and standard errors from three or more independent experiments on astrocytes obtained from separate mice. Representative raw data traces displayed in figures are averaged from at least 3-6 selected ROIs within a single experiment. Parameters measured included amplitude of peak response (change in [Ca^2+^]_i_ from baseline value, actual value or normalized to maximal response), fraction of responding cells (# ROIs with Ca^2+^ change >5% baseline/total # ROIs), time to peak (from >5% baseline change to peak), and area under the curve (AUC: change in Ca^2+^ over time above baseline).

### Drug experiments

To test the role of sphingosine-1-phosphate (S1P) in astrocyte sensitivity to flow, cells were treated for 1-2 hours prior to and for the duration of flow experiments with 1 µM S1P receptor antagonist fingolimod (FTY720) (stock 1 mM in DMSO, Sigma #SML0700). FTY720 experiments were performed with Protocol 1 in flow media supplemented with 0.3% BSA. Conversely, the ability to reverse the nonresponsive phenotype for astrocytes in 0% BSA was tested by >1 hour incubation prior to and for the duration of flow experiments with 1 µM S1P (stock 0.2 mM in 1:5 methanol:water, Sigma #S9666) in 0% BSA flow media using Protocol 1, 10 s. Results were compared to those obtained with vehicle alone (DMSO or methanol).

To test whether the mechanosensitive channel Piezo1 was involved in the responses of astrocytes to flow, we tested effects of an inhibitor (the spider toxin GsMTX 4; Tocris #4912 stock 50 μM in DMSO and the Piezo1 channel activator Yoda1 (Tocris 6586 10 and 20 uM, stock 20 mM in DMSO).

Mediation of astrocyte shear stress sensitivity through the phospholipase C (PLC) pathway was tested through pharmacological inhibition, Cells were treated with 5 µM of the PLC inhibitor U-73122 (MedChemExpress Cat #: HY-13419) after incubation for 1.5 hour prior to and for the duration of flow experiments and tested using Protocols 1.

The role of extracellular glycocalyx in amplification of astrocyte mechanotransduction was tested by degrading chondroitin sulfate proteoglycan (CSPG) with 0.2 U/mL chondroitinase ABC (Millipore Sigma #C2905-2UN) and heparan sulfate proteoglycan with 0.25 U/mL heparanase III (Sigma #H8891). Cells were treated for 1-2 hours at 37°C prior to flow experiments in flow media containing 45 μM BSA and tested using Protocol 1, 10 s, with enzyme omitted from the flow media. Other drugs

| Reagent | Source | Cat # | Concentration |
| --- | --- | --- | --- |
| Bovine Serum Albumin<br>(BSA) | VWR® | 10791-790 | 7.5, 15, 45, 90 µM |
| Sphingosine-1-phosphate | Tocris | 1370 | 1 µM |
| FTY 720 | Tocris | 6176 | 1 µM |
| Chondroitinase | MilliporeSigma | C2905 | 0.2 U/mL |
| Cyclopiazonic acid (CPA) | MilliporeSigma | C-1530 | 10 µM |
| Gadolinium(III) | MilliporeSigma | 439770 | 1 mM, 30 µM |
| Yoda 1 | Tocris | 5586 | 10, 20 µM |
| U-73122 | MCE | HY-13419 | 5 µM |
| GsMTx4 | Tocris | 4912 | 1 µM |
| Heparinase III | MilliporeSigma | H8891 | 0.25 U/mL |
| RN 1734 | Tocris | 3746 | 10, 20, 30 µM |
| HC064074 | Tocris | 4100 | 1, 10 µM |
| GSK2193874 | Tocris | 5106 | 1, 2 µM |
| 4α-Phorbol 12,13-didecanoate (4α-PDD) | MilliporeSigma | P8014 | 1 μM |

### Data analysis

Fluorescence ratios were generally quantified in >20 individual ROIs in each experiment, and values presented in histograms represent means and standard errors from three or more independent experiments on astrocytes obtained from separate mice. Representative raw data traces displayed as figures are averaged from ≥5 selected ROIs within a single experiment. Parameters measured included amplitude of peak response (change in [Ca^2+^] from baseline value, actual value or normalized to maximal response), fraction of responding cells (# ROIs with Ca^2+^ change >5% baseline/total # ROIs), time to peak (from >5% baseline change to peak), and area under the curve (AUC: change in Ca^2+^ over time above baseline).

### Statistics

Multiple ROIs were selected in each field for calculation of Ca^2+^ levels in flow experiments. Mean values obtained in individual experiments were averaged between multiple experiments on different cell cultures to obtain mean and variance values for comparison of amplitudes, number of responding cells and area under curve, as presented. Statistical significance was evaluated using unpaired Student’s t test, One-way ANOVA (Holm-Sidak’s and Tukey’s multiple comparisons test), and two-way ANOVA as described in figure legends using GraphPad Prism 9. Nonlinear Regression was used to calculate EC_50_ using GraphPad Prism 9. p-values are defined as followed: * p<0.05, ** p<0.01, *** p<0.001, **** p<0.0001. All data are presented as mean ± SEM.

## Results

### Properties of astrocyte cell cultures

Purity of astrocyte populations isolated for these experiments was evaluated through immunolabeling of cultures for the astrocyte intermediate filament protein glial fibrillary acidic protein (GFAP) and through imaging mechanical responses from cultures prepared from GFAP::GCaMP6f mouse brains. As illustrated in Fig S1A, most cells exhibited prominent GFAP staining when evaluated at passages 1 and 3 (82.14±1.7 %; 89.3±1.2%); data pooled in histogram from 34 fields of view in Fig S1B).

Additional evidence for preponderance of astrocytes in the cull cultures used for these experiments was obtained from studies in which we measured flow responses of astrocytes isolated from GFAPCre::GCaMP6f mice using BSA-containing flow media. As illustrated in Supplemental movie 1 and in the selected frames shown in Fig S1C-E, virtually all cells showed increased CGaMP6f fluorescence in response to a range of shear flow rates (average of 5 ROIs depicted in Fig S1E shown in Fig S1F-G). Display of Ca^2+^ changes at higher temporal resolution in five selected ROIs (Fig S1 F-G) showed that cells responded to moderate and low flow rates with transient Ca^2+^ increases. Activity patterns in cells within clusters were similar to those of nearby but non-contacting cells (ROIs for selected traces are shown in Fig S1F-G). These measurements indicate that virtually all cultured cells express the GFAP-driven genetically encoded Ca^2+^ indicator GCaMP6, confirming high purity of the cultures used for these studies. Moreover, most cells responded to shear stresses over a wide range of intensities.

### Astrocytes are sensitive to low levels of shear stress

Although GCaMP6f is a bright and highly responsive Ca^2+^ indicator, its fluorescence is monitored at a single wavelength and thus is concentration-dependent and modifiable by cell volume change during swelling or shrinkage. In order to quantify the intracellular Ca^2+^ elevations in astrocytes without this potential confound, we used the ratiometric Ca^2+^ indicator Fura-2, where emission is measured when exciting at two wavelengths straddling the isosbestic point. Wild type primary cortical astrocytes in culture were exposed to calibrated shear stresses in the range of 0 to 20 dyn/cm^2^ for 10 or 30 sec at intervals of 3 min using flow media containing 45 μM BSA. As in the case of GCaMP6 imaging, ratiometric image sequences revealed responses in most cells evoked by shear stress (images taken before, during and after 30 sec shear stresses of 0.19 and 5.78 dyn/cm^2^ are shown in Fig 1B). As illustrated by the display of responses averaged within single experiments assessed with either 10 sec (Fig 1C) or 30 sec (Fig 1D) stimuli, responses to the longer stimulation were larger. For both stimulus durations, magnitude was dependent on stimulus intensity, with responses decreasing at stresses below about 3 dyn/cm^2^ but persisting at even very low flow rates. When normalized Ca^2+^ changes were plotted as a function of shear stress, attenuation was marked below about 1 dyn/cm^2^, with a somewhat lower EC50 for the briefer stimulus duration (Fig 1E). Percentage of total cells responding also revealed strong amplification of sensitivity over the range of shear stress from 0-1 dyn/cm^2^ for both 10 and 30 sec stimuli (Fig 1F). An additional property of the response that was stimulus dependent was the time to peak; for both 10 and 30 sec stimuli, time to peak was rather stable at about 40 sec for stimulus intensities above 2 dyn/cm^2^ but was considerably longer for briefer stimuli (Fig 1G). It is notable that the time to peak greatly exceeded the 10 sec stimulus duration, implying the existence of an intracellular process interposed between stimulus and response in addition to opening of a mechanosensitive channel.

We also examined the responses to stimuli of constant intensity but varied duration. As illustrated in Fig S2A-C, response amplitude more than doubled when duration was extended from 5 to 30 sec, whereas fraction of responding cells was only slightly higher at 30 sec, the longest duration tested, than at 2 sec, the shortest duration.

### Albumin is required for astrocyte high mechanosensitivity to flow

Protein concentration in blood is 6-8 g/dl, of which more than half is serum albumin (>35 g/L or about 0.5-0.8 mM), and although its concentration is much lower in cerebrospinal fluid (CSF), it is still 5 μM or even higher ((Thomson et al., 2018, LeVine, 2016). We examined impact of albumin on astrocyte mechanosensitivity, testing a range of concentrations encompassing those to which astrocyte endfeet are exposed under physiological or pathophysiological conditions. For these experiments, cells were stimulated repeatedly with 10 and 30 sec duration 2.8 dyn/cm^2^ forces, first in normal flow solution (45 μM albumin), then in test albumin concentrations (0-100 μM), followed by stimulation after rinse with normal flow solution (representative images for responses to 0 and 90 μM albumin shown in Fig 2A, measured fluorescence intensities in regions of interest in single experiments for 10 and 30 sec duration stimuli delivered in flow media containing 0 or 45 μM albumin shown in Fig 2D). When normal flow solution (containing 45 μM BSA) was exchanged for solution containing 15 μM or higher albumin concentration, astrocyte Ca^2+^ responses to either 10 or 30 sec duration stimuli did not change in amplitude (Fig 2B) or in fraction of responding cells (Fig 2E), indicating that 15 μM or slightly lower is the concentration at which the response is maximal. Responses to lower concentrations of albumin were smaller in amplitude (Figure 2B-D), and in the absence of albumin the response to shear stress stimulation was almost completely abrogated (shown in each panel of Fig 2; note that when area under the curve for responses at 30 sec were computed, the difference in responses was more than ten-fold). EC50 values calculated for amplitudes of responses to constant shear intensity and duration in different albumin concentrations ranged from 3-10 μM (Fig 2B) and from about 10 to 50 μM when fraction of responding cells was quantified (Fig 2E). Thus, astrocyte sensitivity to flow is vastly amplified by albumin, with EC50 in the range from 0.3-1 μM. As discussed, it is remarkable that this set point is within the range of protein concentrations that is present in normal CSF (20-60 mg/%(Thomson et al., 2018)).

### S1P receptors mediate amplification of astrocyte Calcium signaling in response to shear stress

Shear stress responses of endothelial, smooth muscle and other cell types is highly dependent on the presence of serum albumin, an effect that has been attributed to direct maintenance of the cell’s glycocalyx and more recently to the presence of sphingosine 1-phosphate (S1P) carried by the serum protein (Zhang et al., 2016, Tarbell et al., 2014). To test whether albumin sensitization of astrocyte responsiveness is also dependent on S1P, we evaluated whether response in absence of serum was restored by addition of S1P to BSA-free flow medium and whether sensitivity was reduced in the presence of S1P receptor inhibition.

To test directly for an effect of S1P on astrocyte mechanosensitivity, we treated cells with 1 µM S1P for at least 1 hour in the absence of albumin, a condition under which astrocytes are normally unresponsive (see Fig 2). Cells were exposed to flow media lacking BSA, to which they did not respond, and then S1P was added and shear stress was reapplied. As shown in image frames in Figure 3A taken before, during and after stimulation at 5.8 dyn/cm^2^, responses were absent in flow medium lacking BSA but were restored in the presence of BSA. Amplitudes of responses and number of responding cells were both analyzed to quantify the impact of S1P on stimulus-response relations, which demonstrated a clear restoration of mechanosensitivity relative to 0% BSA vehicle control. In the presence of S1P shear rate dependence of Ca^2+^ peak amplitude (EC50 = 1.23 dyn/cm^2^, Fig 3B) and percentage of responding cells (EC50 = 0.21, Fig 3E) were consistent with responses obtained in the presence of BSA (compare Fig 2). Overall responsiveness, evaluated by calculation of area under the curve in response to maximal stimulus (5.8 dyn/cm^2^, 10 sec), was similar to that obtained in the presence of 45μM BSA (compare Fig 3C with Fig 2C).

**Figure 3:**
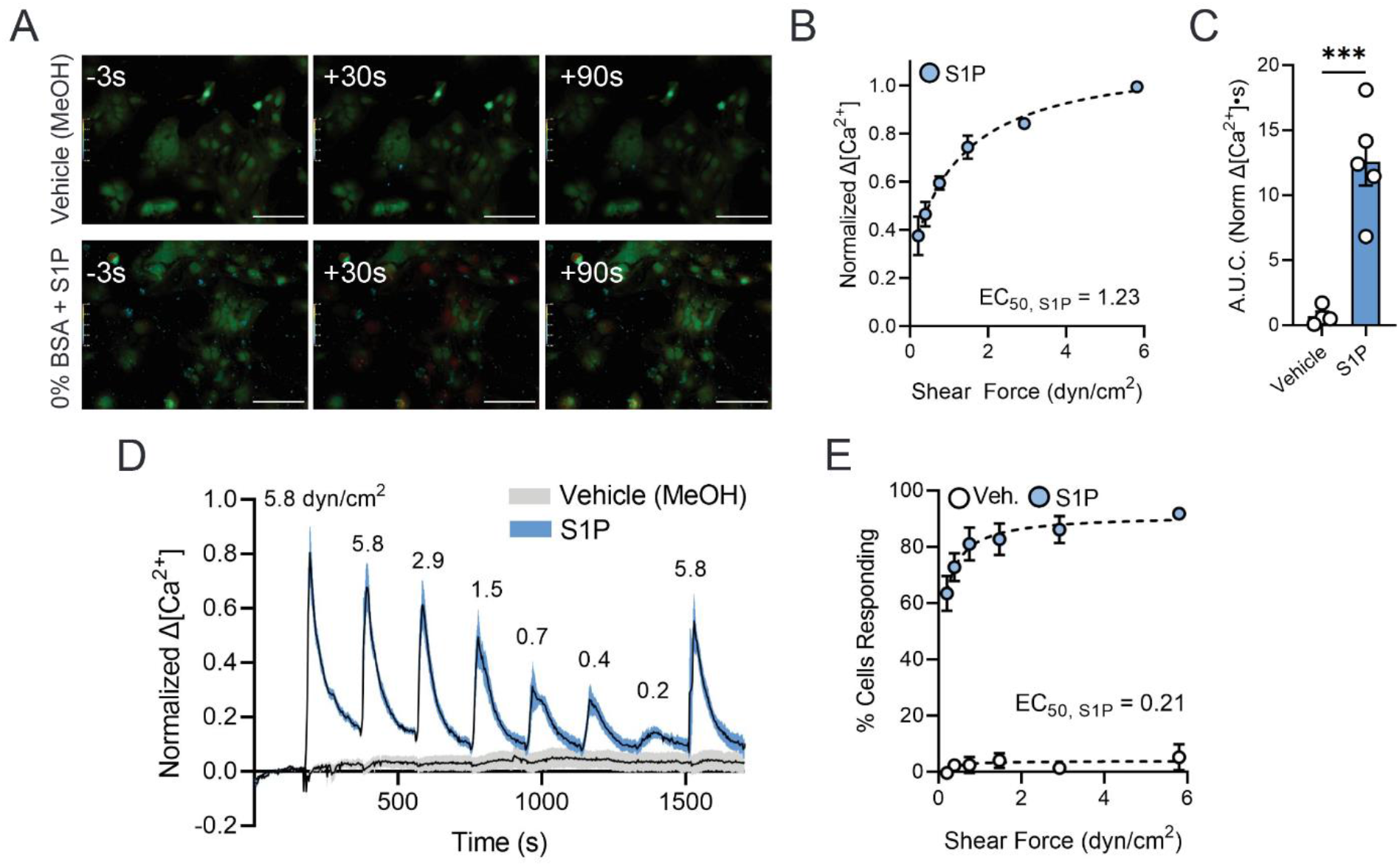
S1P receptor antagonist, FTY720, blunts astrocyte response to shear stress. (A) Representative ratiometric fluorescence micrographs of Ca^2+^ response time course in astrocytes in 45 BSA plus vehicle (DMSO) and 45 BSA plus FTY720 in response to 10 sec 2.9 dyn/cm^2^ shear. Scale bars 100 µm. (B) Normalized change in Ca^2+^ concentration amplitude across shear rates with 10s stimuli. (C) Area under normalized Ca^2+^ response curve for vehicle control and FTY720 conditions at 2.9 dyn/cm^2^. Unpaired Student’s t test: *p < 0.05. (D) Representative trace of change in Ca^2+^ concentration in 45 BSA plus FTY720 normalized to DMSO control. (E) Percentage of cells responding to a range of shear rates for 10s stimuli. All data shown as mean ± SEM. n ≥ 3 for all experiments.

To evaluate the effect of inhibiting S1P receptor activity, we applied the broad spectrum S1P receptor modulator fingolimod (FTY720). After incubation with FTY720 for 1 hr, astrocyte responses to flow were substantially blunted (Fig 4A). The decreased response amplitude was evident across the entire range of stimulus intensities (Fig 4B, D), with only slightly reduced EC50 values compared to vehicle control. Area under the curve in response to 2.8 dyn/cm^2^ was also significantly reduced by FTY720 (Fig 4C, as was the % responding cells at all stimulus intensities (Fig 4E)).

**Figure 4:**
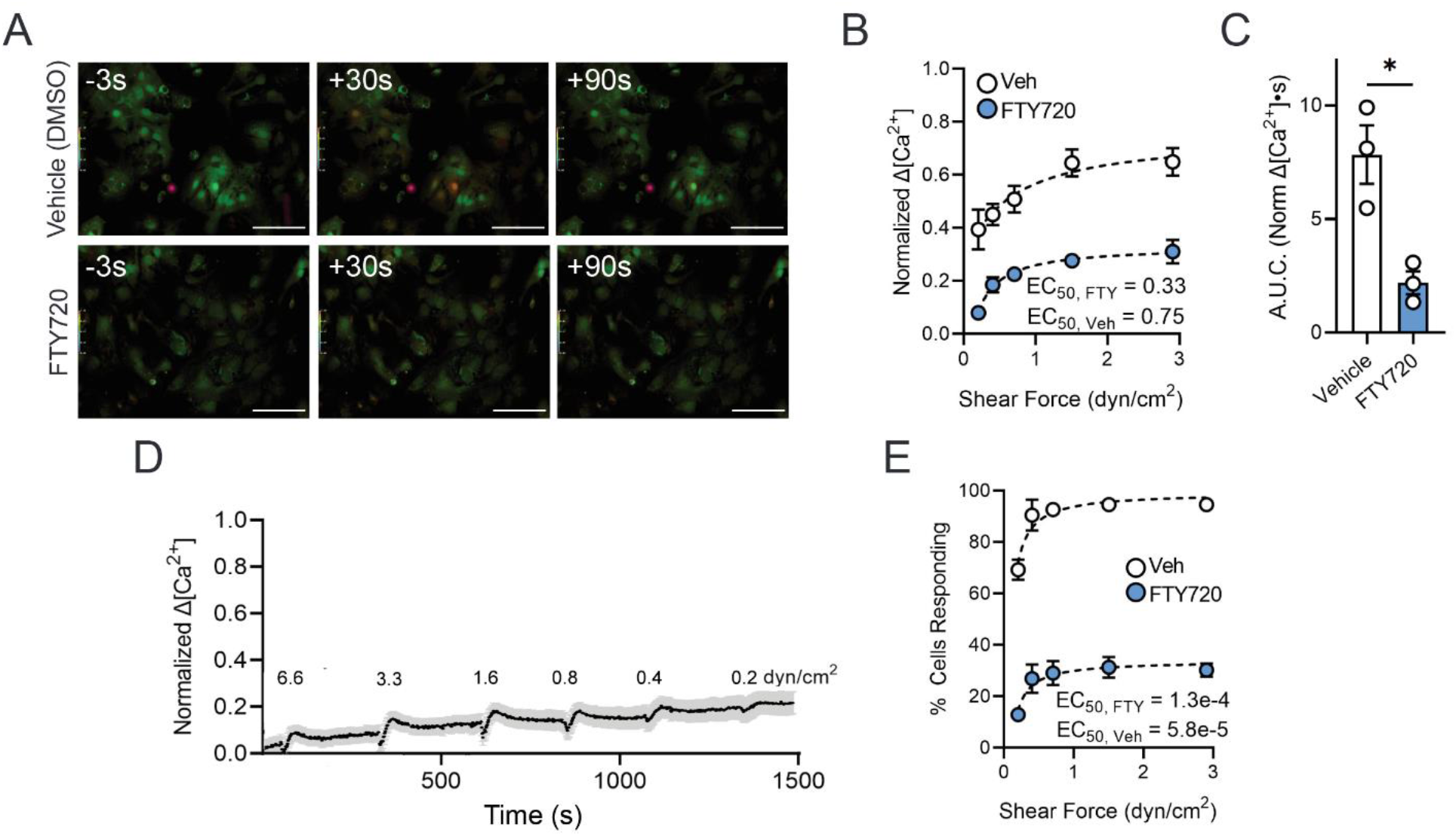
S1P recovers astrocyte mechanosensitivity in albumin-free condition. (A) Representative ratiometric fluorescence micrographs of Ca^2+^ response time course in astrocytes in 0 BSA plus vehicle (methanol) and 0 BSA plus S1P across a range of shear stresses. Scale bars 100 µm. (B) Normalized change in Ca^2+^ concentration amplitude across shear rates with 10s stimuli. (C) Area under normalized Ca^2+^ response curve for vehicle control and S1P conditions at 5.8 dyn/cm^2^. Unpaired Student’s t test: ***p < 0.001. (D) Representative traces of normalized change in Ca^2+^ concentration in 0 BSA with vehicle or S1P. (E) Percentage of cells responding to a range of shear rates for 10s stimuli. All data shown as mean ± SEM. n ≥ 3 for all experiments.

### Piezo1 is the target of S1P-amplified astrocyte shear stress sensitivity

To identify the mechanosensitive channel underlying astrocyte shear stress sensitivity, we tested effects of pharmacological inhibitors on astrocyte responses. These compounds, their presumed molecular targets and tested concentrations are listed in Table 1. As shown in Fig S4 and S5, several inhibitors reduced response amplitude by >20% suggesting that they may participate in astrocyte shear stress responses. Among the most potent of the inhibitors tested was the tarantula toxin, GsMTx-4, which produced a blockade that was progressive as subsequent stimuli were applied following its wash-in and was gradually restored upon washout (representative experiment in Fig 5A, summary of results in Fig 5B). To further test the involvement of Piezo1 in the astrocyte shear stress responses, we applied Yoda1, an agonist that enhances Piezo1 activity. In these experiments, we first conducted a series of shear stress stimulations under conditions of 0, 7.5 and 45 μM BSA, and then washed in Yoda1 and repeated the series of shear stresses. As shown in typical experiments for 0 and 45 μM BSA in Figs 5C, D, and quantified from all experiments in Fig 5E, addition of Yoda1 strongly potentiated the response to shear stress, even enabling responses at all flow levels that were normally absent in solution lacking BSA (Figs 5C, E). To test the hypothesis that Yoda1 was acting to enhance activation of Piezo1 through S1P receptors, we repeated experiments in the presence of FTY720. As illustrated in Fig 5F and quantified in Fig 5G, shear stress responses were absent in the presence of FTY720 but responses to even very small shear stresses were rescued by treatment with Yoda1. The amplification of shear stress sensitivity by Yoda1 provides strong evidence that Piezo1 contributes substantially to the observed mechanosensitivity.

**Figure 5:**
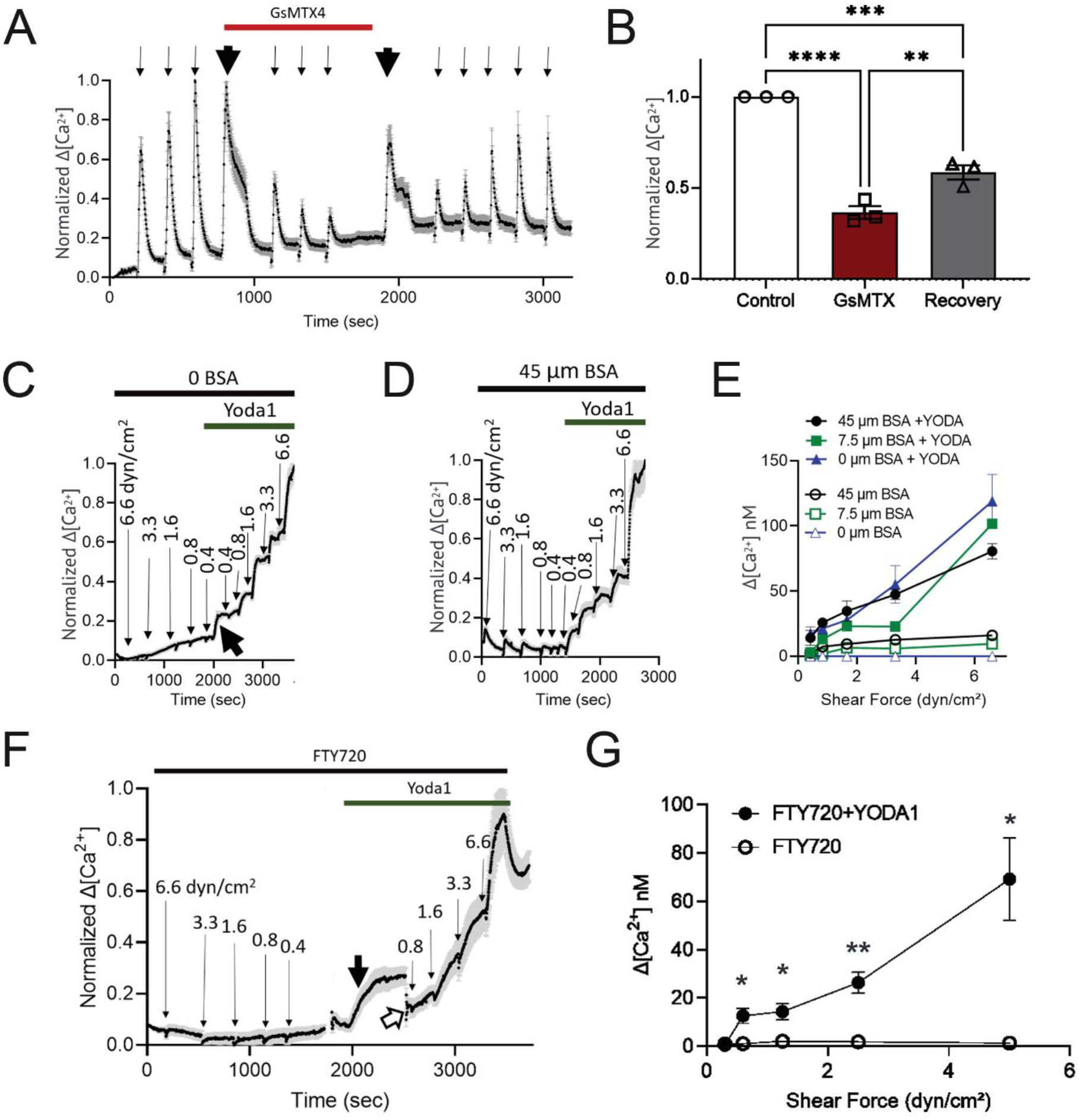
PIEZO1 determines the mechanosensitivity of astrocytes. (A) Representative experiment showing that GsMTx4 (1 µM), a PIEZO1 antagonist, reversibly inhibits the astrocyte calcium responses to flow. 5.8 dyn/cm^2^ shear flow for 10 seconds (thin arrows) was applied at 300-second intervals before, after drug additions and after wash-out. Wash-in and wash-out of drug (thick arrows) were done at 0.2 ml/min for 2 minutes. (B) Summary of 3 independent experiments showing the reduction of shear-induced response by 1 µM GsMTX4 (red bar) and partial recovery (gray bar) upon washout. One-way ANOVA followed by Tukey’s post hoc test: **p<0.01; ***p<0.001; ****p<0.0001. (C, D) Representative experiments showing the amplification of astrocyte calcium shear response by Yoda1, a PIEZO1 activator. A descending series of 10-second shear forces was applied in flow medium containing 0 or 45 µM BSA (C and D respectively), then 10 µM Yoda1 was added and an ascending series of 10-second shear forces was applied. Shear forces (dyn/cm^2^) are indicated above each response. Note: substantially enhanced responses in the presence of Yoda1. (E) Summary data comparing Yoda1 effect in various BSA concentrations (0, 7.5, and 45 µM) at moderate shear forces. Note: significant enhancement of sensitivity in the presence of Yoda1 at all BSA concentrations. Two-way mixed ANOVA: 45 µM BSA with vs without Yoda: p-value = 0.0024; F(1,13) = 14.15; 7.5 µM BSA with vs without Yoda: p-value = 0.0021; F(1,21) = 12.28; 0 µM BSA with vs without Yoda: p-value = 0.0202; F(4,28) = 3.468. (F) Representative experiment of calcium response of astrocytes incubated in FTY720 for 1.5 hours. A descending series of 10-second shear forces was applied in flow medium containing 45 µM BSA, then 10 µM Yoda1 was added (bold black arrow) and an ascending series of shear forces was applied; open arrowhead indicates refocusing. (G) Summary of 3 independent experiments showing the block of flow-induced stress after FTY720 incubation and the recovery of response upon the addition of Yoda1. Unpaired Student’s t test: *p<0.05; **p<0.01. n ≥ 3 for all experiments. All data shown as mean ± SEM.

### Inhibition of phospholipase C (PLC), a downstream target of S1PR activation, eliminates astrocyte shear stress induced Ca^2+^ signaling

Blockade of response by fingolimod implicates G-Protein coupled signaling through S1P receptors. Fingolimod blocks S1PR 1, 3, 4 and 5, which signal through G-proteins to activate PLC, Ras, ROCK, and PI3 kinase. With the exception of the PLC inhibitor U-73122, drugs tested produced either slight or inconsistent inhibitions of astrocyte responses to flow. After incubation in the PLC inhibitor for 1.5 hour, however, astrocytes failed to respond to the full range of stimulus intensities (Fig 6C). In order to determine whether the effect of U-73122 was through the action on Piezo1 channels, we re-evaluated the responses following chemical activation of Piezo1 by Yoda1. As shown in Fig 6B and C, responsiveness was absent in the presence of the PLC inhibitor but was rescued by treatment with Yoda1.

**Figure 6:**
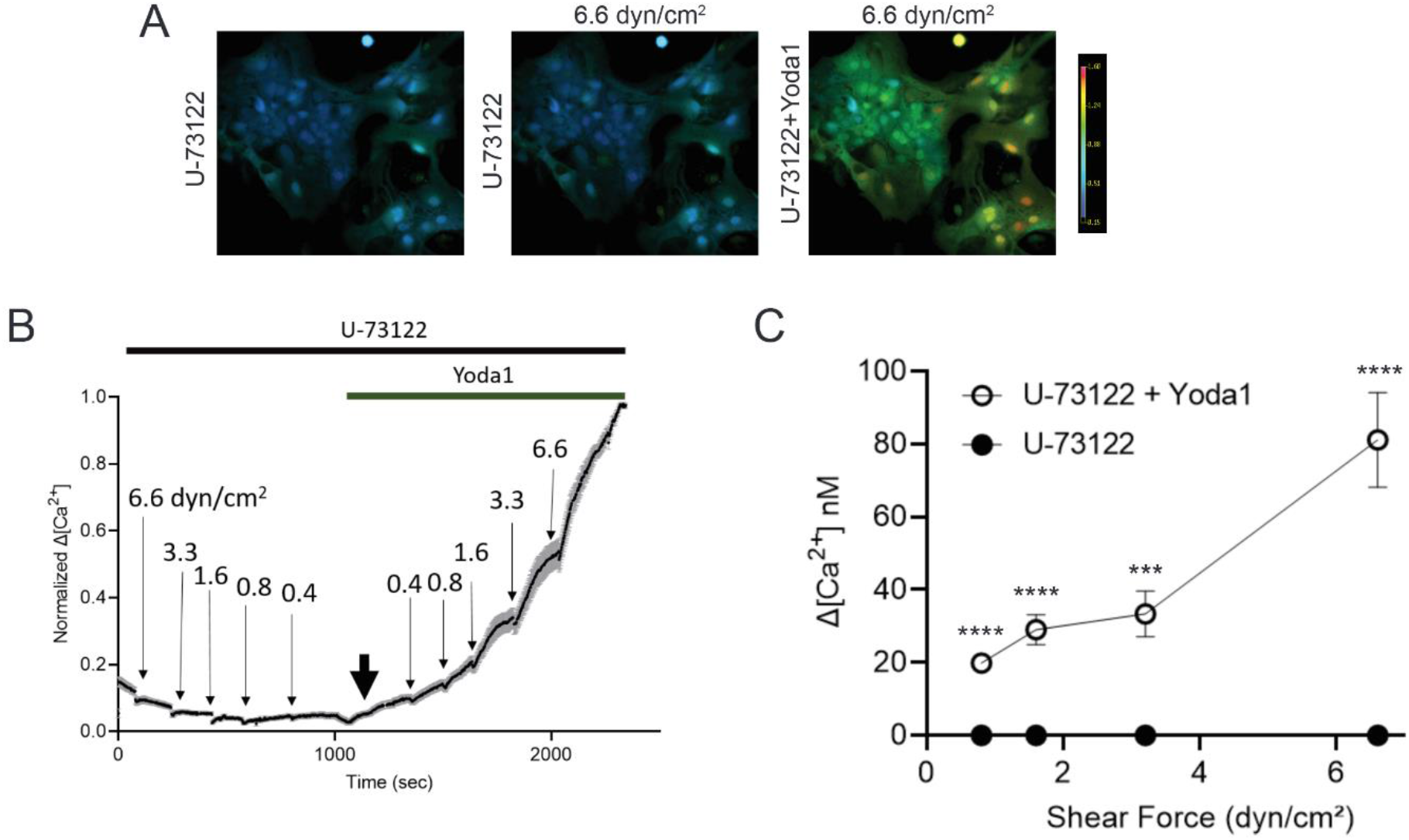
PLC inhibition eliminates astrocyte Ca^2+^ response. (A) Representative ratiometric fluorescence micrographs of Ca^2+^ response time course in astrocytes in 45 µM BSA plus PLC inhibitor, U-73122, across a range of shear rates. Scale bars 100 µm. (B) Representative traces of normalized change in Ca^2+^ concentration showing the blockade of shear response after incubation in U-73122 for 1.5 hr. A descending series of 10-second shear forces was applied in flow medium containing 45 µM BSA and U-73122, then 10 µM Yoda1 was added (bold black arrow) and an ascending series of shear forces was applied. (C) Summary of 6 independent experiments showing the block of flow-induced stress after U-73122 incubation and the recovery of response upon the addition of Yoda1. Unpaired Student’s t test: ***p<0.001; ****p<0.0001. All data shown as mean ± SEM.

### Extracellular proteoglycan matrix participates in astrocyte mechanosensitivity

Astrocytes secrete and are attached to proteoglycans and glyosaminoglycans that fill interstitial space in the brain and provide a reservoir for lipophilic molecules, including S1P ((thi et al., 2013a, Zeng et al., 2018). Previous studies on endothelial and smooth muscle cells showed that shear stress responses in those cell types are greatly attenuated by enzymatic removal of the glycocalyx (Ebong et al., 2014a). In order to determine whether these extracellular space molecules play a role in astrocyte shear responses, we immunostained astrocyte cultures for three major components of the brain extracellular space: chondroitin sulfate proteoglycans, laminin, and heparan sulfate proteoglycans. We then treated the cells with enzymes to degrade the proteoglycans and tested the consequence for astrocyte mechanosenstivity. As shown in figure 7 and Fig S6, we found abundant surface expression of these extracellular matrix components on the astrocytes. For HSPG, treatment for 2 hr with heparinase III decreased the expression by 53 ± 3.5 %. Treatment with chondriotinase for 2 hours produced a similar degree of loss of CSPG (Fig. S6B-D). As shown in Fig 7F, treatment with heparanase dramatically attenuated response to all intensities of shear stress, whereas CSPG cleavage had minimal effect on the responsiveness.

**Fig 7:**
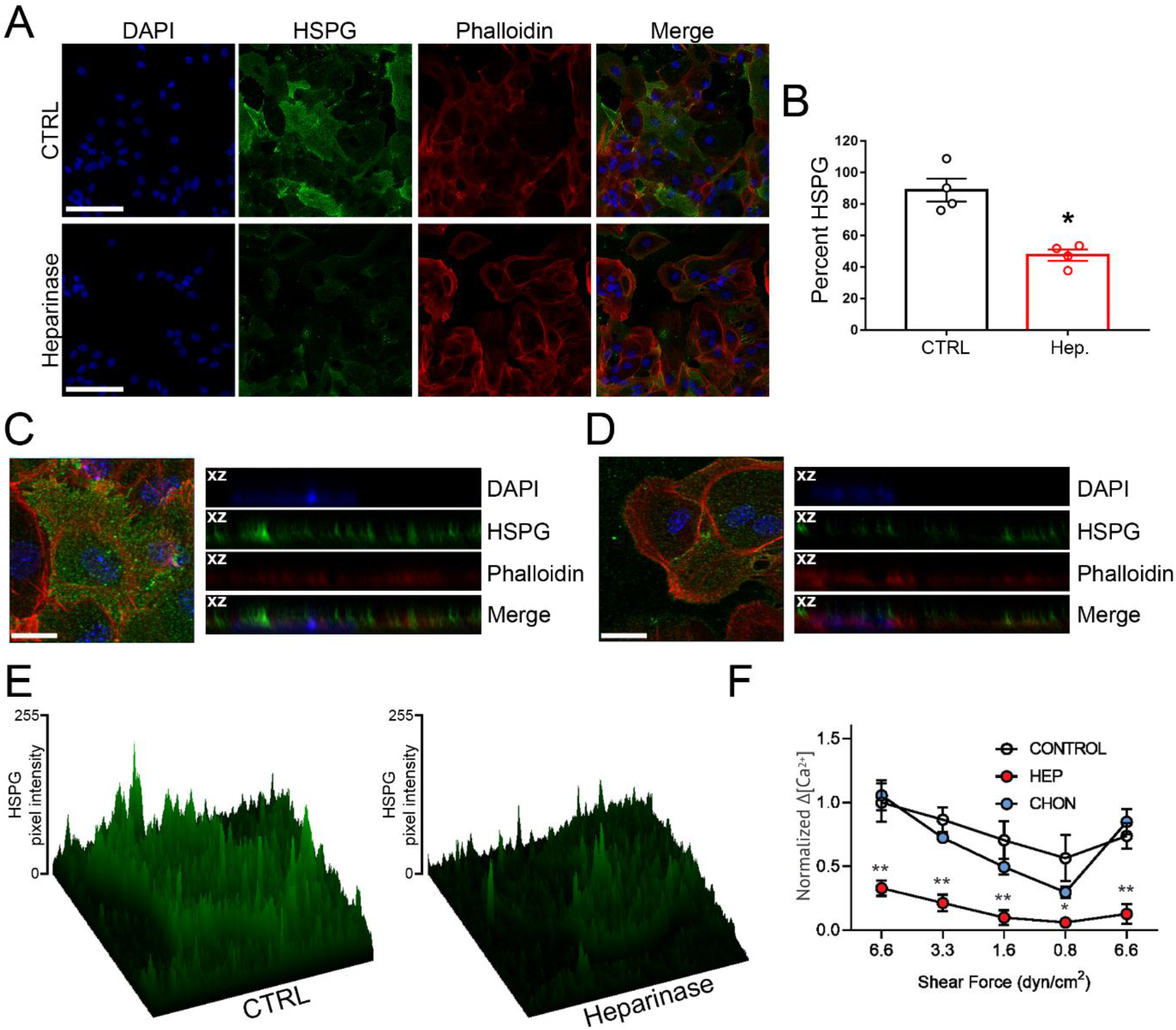
Attenuation of responses by heparan sulfate proteoglycan degradation. (A) Representative single-slice confocal images of HSPG (green), F-actin cytoskeleton visualized with phalloidin (red) and their merge in WT astrocytes. Cells were cultured in DMEM without (CTRL) and with heparinase III (Heparinase) for 2 h. The nuclei were stained blue with DAPI. Scale bar, 50 μm. (B) Histogram showing HSPG fluorescence intensity quantification expressed as a percentage relative to that of the control (CTRL, black), considered as 100 % after 2 h treatment with heparanase III (Hep., red). Data are expressed as mean ± SEM. Four images per experiment, from a total of four experiments, were taken for each condition. Unpaired Student’s t test: *p < 0.05. (C) Left, confocal maximum intensity projections of control WT astrocytes immunostained for HSPG (green), F-actin (red) and nuclei (blue). Scale bar, 20 μm. Right, xz cross-sectional views of the stacked confocal images from the right showing the degree of cell surface distribution between HSPG (green), actin cytoskeleton (red) and nuclei (blue). (D) Left, confocal maximum intensity projections of WT astrocytes after 2 h treatment with heparanase immunostained for HSPG (green), F-actin (red) and nuclei (blue). Scale bar, 20 μm. Right, xz cross-sectional views of the stacked confocal images from the right showing the degree of cell surface distribution between HSPG (green), actin cytoskeleton (red) and nuclei (blue). (E) Reconstructed 3D surface plot images of (C) and (D). Color range from black to green indicates the relative level of fluorescence intensity in pixel. (F) Summary of normalized astrocyte calcium responses to a range of shear forces before and after enzymes degrading heparan sulfate proteoglycan or chondroitin sulfate proteoglycan. n=3, 4, 8 for control, HSPG degradation, CSPG degradation, respectively. Unpaired Student’s t test: *p<0.05; **p<0.01. All data shown as mean ± SEM.

## Discussion and Conclusions

### Forces that the endfoot feels

Astrocytes are exposed to diverse mechanical forces, including rhythmic pressure differentials in CSF driven by the cardiac cycle, traumatic forces in concussive brain injury or multiple mild impacts as in soccer heading, interactions with substrates of varying stiffness, as in glial scars, as well as membrane stretch due to osmotic volume changes and stretch due to vasomotion. Previous studies have measured astrocyte Ca^2+^ responses to focal pressure pulses (Turovsky, Braga et al. 2020), pressures across the arteriolar wall (Kim and Filosa 2012, Kim, Iddings et al. 2015) and brief high intensity concussive forces (Ravin, Blank et al. 2012, Maneshi, Sachs et al. 2015, Ravin, Blank et al. 2016, Maneshi, Maki et al. 2017, Maneshi, Sachs et al. 2018, Ravin, Morgan et al. 2019). In each of these studies, sensitivity was quite low.

However, astrocyte endfeet form a cylindrical perivascular boundary in which shear forces would be expected to be generated by flow of the perivascular glymphatic circulation (Iliff et al., 2012) along the vascular cells. Thus, even low flow through the perivascular space should generate shear stresses to which astrocytes may respond. Consistent with this hypothesis, we report here (Fig 1) that astrocytes respond to shear stresses well below 0.1 dyn/cm^2^ when measured using solutions approximating the composition of cerebral spinal fluid. To put this finding into context, flow velocity measured from movements of injected nanoparticles in perivascular space (Min Rivas et al., 2020) permits calculation of shear stress in this compartment; from Fig. 4C of that paper, dV/dt is about 12/sec and shear stress at the wall is about 0.12 dyn/cm^2^. [Note that this calculation assumes that the perivascular space is empty; shear stress would be much higher if matrix were present and impeded flow.] We conclude that astrocytes are very responsive to mechanical force magnitudes to which their endfeet are normally exposed.

Other recent studies provide additional evidence that low level mechanical stimuli in brain can have physiological effects. For example, treadmill running and passive head movements in rodents were reported to exert interstitial shear stress of 10-30 dyn/cm^2^ in prefrontal cortex (Ryu et al., 2020), a force sufficient to internalize neuronal serotonin receptors. Low shear stress with protein-containing solution applied to cultured astrocytes (0.5 dyn/cm^2^) was shown to greatly enhance phagocytosis of lysed cells (Wakida et al., 2020). CSF shunt implantation is common treatment for hydroceophalus, often failing due to development of astrocyte scar at regions of shear stress >0.5 dyn/cm^2^ (Lin et al., 2003). Astrocytes in 2% FBS responded to 0.5 but not 0.05 dyn/cm^2^ shear stress with cytokine release (Khodadadei et al., 2021). Moreover, gene expression in astrocytes is modified by both pressure driven fluid shear stress and by electro-osmosis due to direct current as used in transcranial stimulation (Cancel et al., 2022).

### Enhancement of response by albumin

The shear stress sensitivity that we measured in astrocytes is much higher than that reported in most previous studies. One likely methodological difference is the use of simple saline solutions lacking protein in previous flow studies. Extensive studies of endothelial and osteocyte responses to shear stress, including our own (Ebong et al., 2014b, Lopez-Quintero et al., 2013, Thi et al., 2004), have shown that shear response depends upon inclusion of albumin in flow medium, which is hypothesized to minimize glycocalyx shedding and maximize availability of lipid effectors (Zeng et al., 2018). With regard to albumin concentration, it is important to note that although protein levels within CSF are vastly lower (by 99%) than in plasma, the levels found to amplify responses in our studies (EC50 3-9 μM) are within the critical normal to pathological range. A levels of CSF protein, consisting mostly of albumin, are about 8 μM in infants, about 15 μM in adults (Thomson et al., 2018), and much higher following infarct or in conditions such as *neuromyelitis optica* and multiple sclerosis (LeVine, 2016). Our studies indicate that shear stress sensitivity is modulated over this normal range of albumin concentrations, and maximal sensitivity occurs at levels seen following immune challenge.

Previous studies beginning more than twenty years ago revealed that astrocytes responded with Ca^2+^ elevation when exposed to albumin, and that this response was abrogated by depletion of lipoproteins in the albumin (Nadal et al., 1995, Nadal et al., 1997). Recent studies on endothelial cells have shown that sphingosine-1-phosphate (S1P) preserved the shear stress responses that were otherwise abolished in the absence of albumin and that inhibition of S1P receptors by FTY720 (fingolimod) blunted mechanosensitivity in the presence of serum protein (Mensah et al., 2017), (Zhang et al., 2016). We have found very similar effects on astrocyte responses to shear stress: S1P restored the blunted sensitivity in absence of albumin (Fig. 3) and treatment with the antagonist largely blunted shear responses in the presence of albumin (Fig. 4).

To determine whether the amplification of the shear responses by S1P was mediated through S1P receptors, we applied the broad-spectrum antagonist FTY720 (fingolimod), which greatly attenuated response amplitude at all stimulus intensities. These findings suggest that the potentiation of astrocyte mechanoresponsiveness by CSF albumin levels results in part from S1P, raising the questions of which downstream pathway is responsible and how albumin mediates S1P activation. Pharmacological experiments using the selective PLC inhibitor U-73122 achieved striking response blockade, implicating this pathway in the response.

### Role of extracellular matrix

Brain extracellular matrix of contains relatively high amounts of glycosaminoglycans (including both chondroitin sulfate and heparin sulfate bound to protein) and relatively low amounts of collagens and fibronectin (Novak and Kaye, 2000). We have reported that cultured astrocytes possess a glycocalyx consisting primarily of heparin sulfate proteoglycans linked to actin cytoskeleton (thi et al., 2013a); in the same report, treatment with heparanase altered endothelial cell cytoskeleton. Our finding in this study that heparanase cleaved the glycocalyx and proportionately decreased mechano-responsiveness is consistent both with the hypothesis that the glycocalyx plays a direct role in mechanotransduction and that it acts as a local reservoir for S1P and perhaps other lipid mediators.

### Role of Piezo1

Albumin in the perfusion solution, and presumably S1P in the albumin, greatly amplifies the astrocyte sensitivity to shear stress. This amplification is likely due to modulation of the shear stress activated mechanosensitive channels. To identify which mechanosensor is required for shear transduction, we tested pharmacological inhibitors of known mechanoreceptors, namely the multimodal TRPV4 channel and the true mechanoreceptor Piezo1. Responses were slightly reduced by one of three TRPV4 receptor antagonists tested, suggesting possible contribution to the responses. By contrast, the tarantula neurotoxin GsMTx4, a moderately selective Piezo1 inhibitor, rapidly and effectively blocked mechanical responses in astrocytes. Furthermore, the chemical Piezo1 activator Yoda1 greatly enhanced response sensitivity in both the presence and absence of albumin and in the presence of FTY720, conditions in which responses are normally absent. As drawn in the graphical abstract, we conclude from these studies that albumin in CSF contains S1P that binds to its receptor to activate phospholipase C. PLC then amplifies the response of Piezo1 to shear stress, leading to elevated intracellular Ca^2+^ and other downstream responses.

Constitutive sensitization of PIEZO1 by S1PR signaling sets the threshold for response to glymphatic circulation. It is notable that the set point for Piezo1 activation is shear force less than 0.1 dyn/cm^2^ in cells exposed to ≥ 8 μm albumin, values that are consistent with measured perivascular flow and normal CSF albumin concentration, as noted above. Even very small variations in either force or CSF protein profoundly affects astrocyte responses. The rapidity by which protein removal blocks astrocyte response indicates that the set point for PIEZO1 gating is constitutively maintained, and total inhibition strongly implicates the involvement of the downstream PLC pathway. That S1PR signaling sets basal activity of ion channels in various cell types has been shown previously, such as in stimulating vascular development through Src kinase stimulation of baseline Piezo1 activation (Kang et al., 2019) and in mediating nociceptor sensitivity through regulation of KCNQ excitability by S1P (Hill et al., 2018). However, this is the first report that S1PR signaling is involved in the astrocyte mechanosome and that its downstream pathway generates constitutive responsiveness of Piezo1.

## Acknowledgements

These studies were facilitated by initial support of DB-G through a collaboration between Profs. Enrico Nasi and Pilar Gomez, Universidad Nacional de Colombia, and the Department of Neuroscience at Einstein and were supported by NIH grants NS092466 and NS116892 to DCS. This research was also supported in part through the Rose F. Kennedy Intellectual and Developmental Disabilities Research Center (IDDRC), which is funded through a Eunice Kennedy Shriver National Institute of Child Health & Human Development center grant (NICHD U54 HD090260-06). Confocal microscopy was supported by Shared Instrumentation grants to the Analytical Imaging Facility. We gratefully acknowledge assistance in related experiments and analysis by Lily Kolb (currently MD student at Donald and Barbara Zucker School of Medicine at Hofstra/Northwell). We also acknowledge highly helpful discussions with Dr. Praveen Ballabh (Einstein/Montefiore) regarding cerebrospinal fluid composition and serum sensitivity of astrocytes and Dr. John Tarbell (CCNY) regarding shear stress in perivascular and parencyhymal spaces.

## Contributions

Experimental design: DD-B, SM, GLY, AC, DCS; performing experiments and data analysis: all; written document: DD-B, SM, GLY, AC, DCS

## SUPPLEMENTAL INFORMATION

**Supplemental Figure 1:**
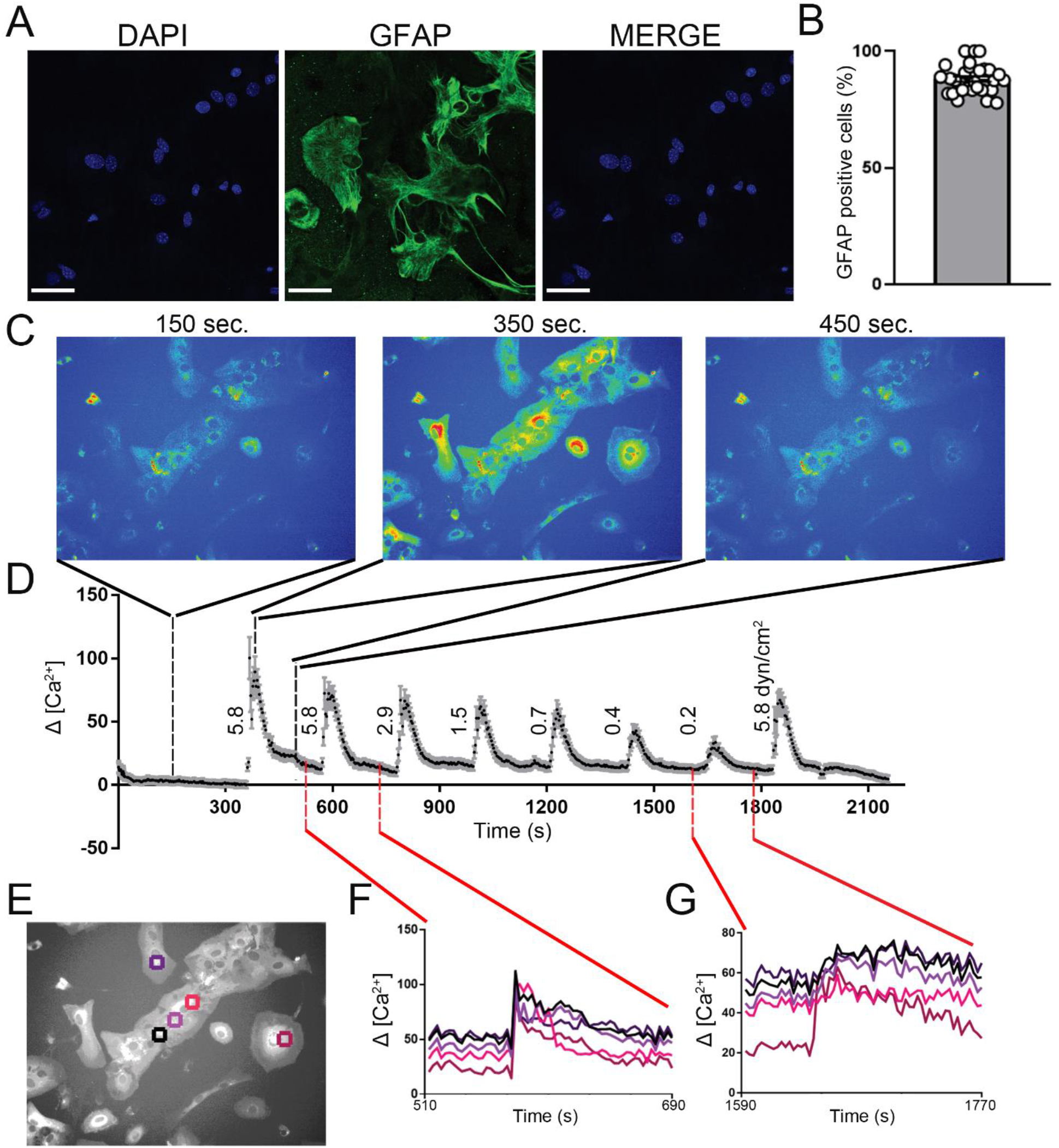
Primary astrocytes cultured in flow chambers are highly homogeneous as judged by GFAP expression in WT and Ca^2+^ responses in GFAPCre::GCaMP6f cells. (A) Confocal micrographs showing GFAP immunostaining; GFAP shown in green and DAPI shown in blue. Scale bars 50 µm. (B) Percentage of cells in cultures stained positive for GFAP at passages 1 and 3 (data pooled from 34 regions of interest (ROIs) in 4 culture dishes; displayed as mean ± SEM. (C) Frames from pseudo-colored GCaMP imaging experiment (where blue indicates lowest and red highest fluorescence levels, which are proportional to intracellular calcium levels in these GFAPCre::GCaMP6f astrocytes) showing average changes occurring before, during and after application of 30 sec 5.8 dyn/cm^2^ stimuli. Graph in D represents averaged values within ROIs throughout the entire field of view in response to a series of 30 sec exposures to shear force magnitudes indicated alongside the responses, ranging from 5.8 to 0.2 dyn/cm^2^. (E) Regions of interest (ROIs) showing responses of adjacent and nonadjacent astrocytes to flow; traces of cell responses to high and low magnitude stimuli correspond to colors of ROIs.

**Supplemental Figure 2:**
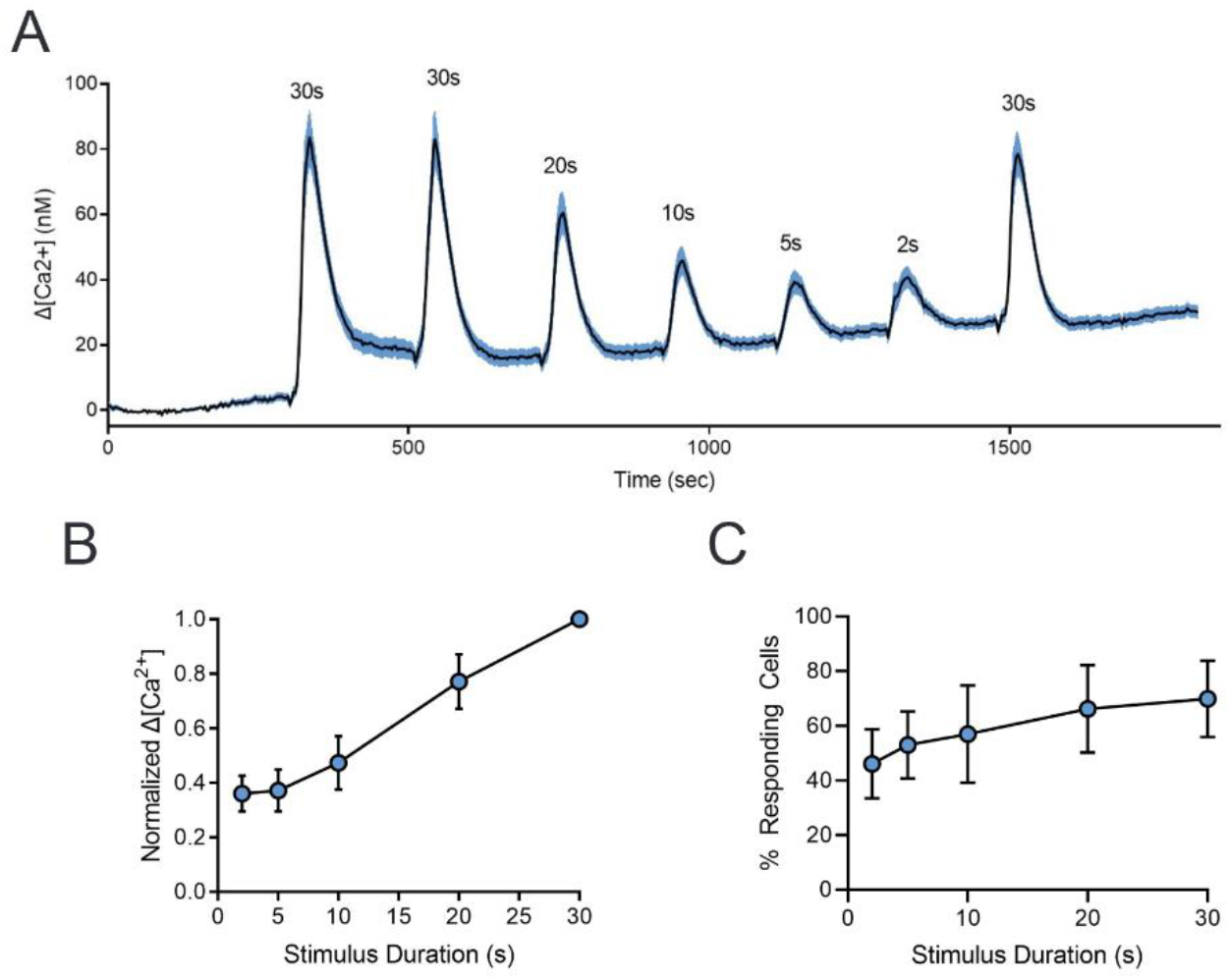
Astrocyte shear stress response is dependent on duration of stimulus. (A) Representative trace of Ca2+ response over a range of 5.8 dyn/cm2 stimulus durations. (B) Amplitude of calcium response as a function of stimulus duration. (C) Percentage of responding astrocytes as a function of stimulus duration. All data shown as mean ± SEM. n = 3 for all experiments.

**Supplemental Figure 3:**
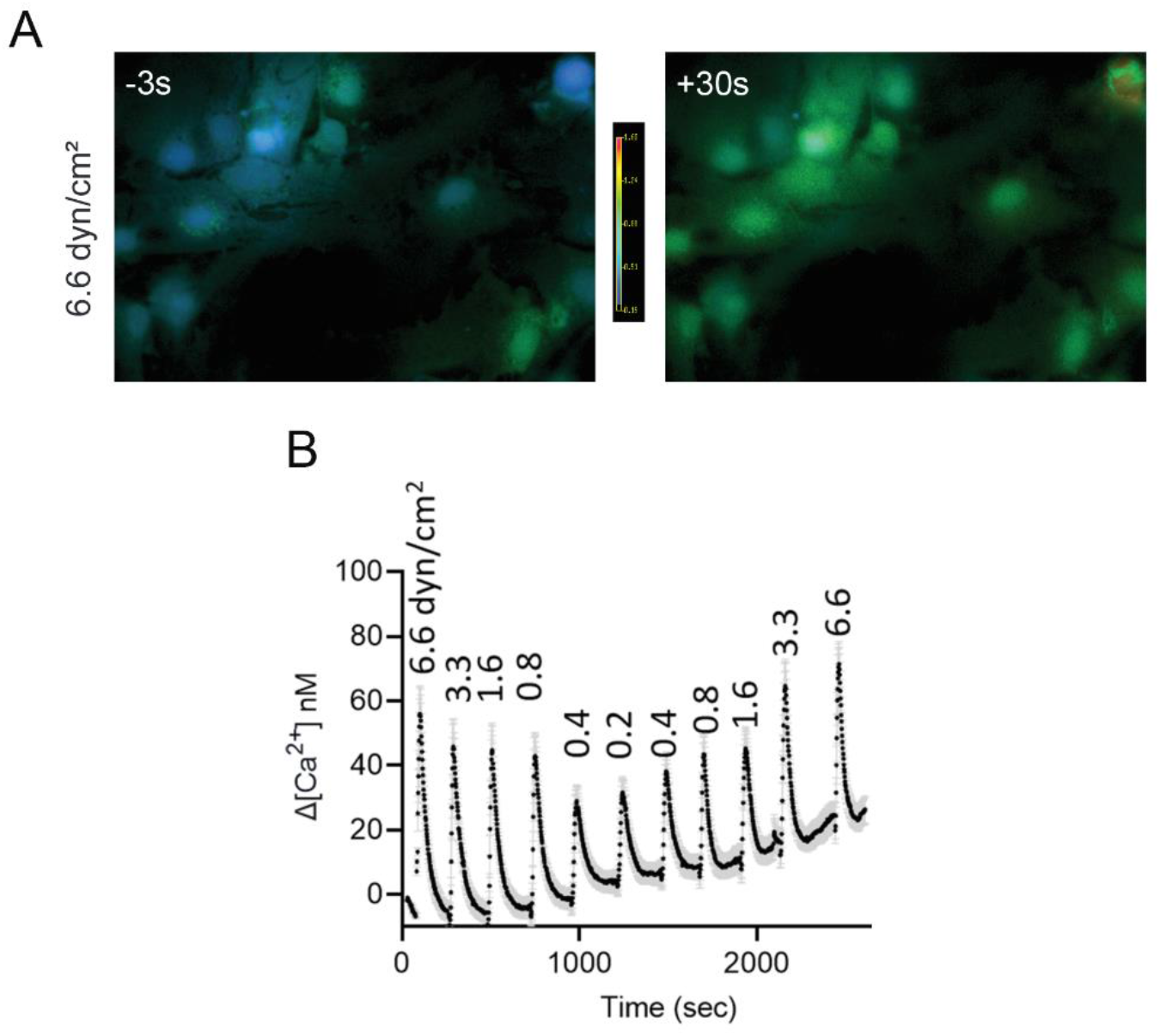
Astrocyte shear stress response is independent of the order by which stimulus is applied. (A) Representative ratiometric fluorescent micrographs of Ca2+ response time course in astrocytes at high (6.6 dyn/cm2) shear forces. (B) Representative trace of Ca2+ response over a range of 6.6 dyn/cm2 stimulus durations.

**Supplemental Figure 4:**
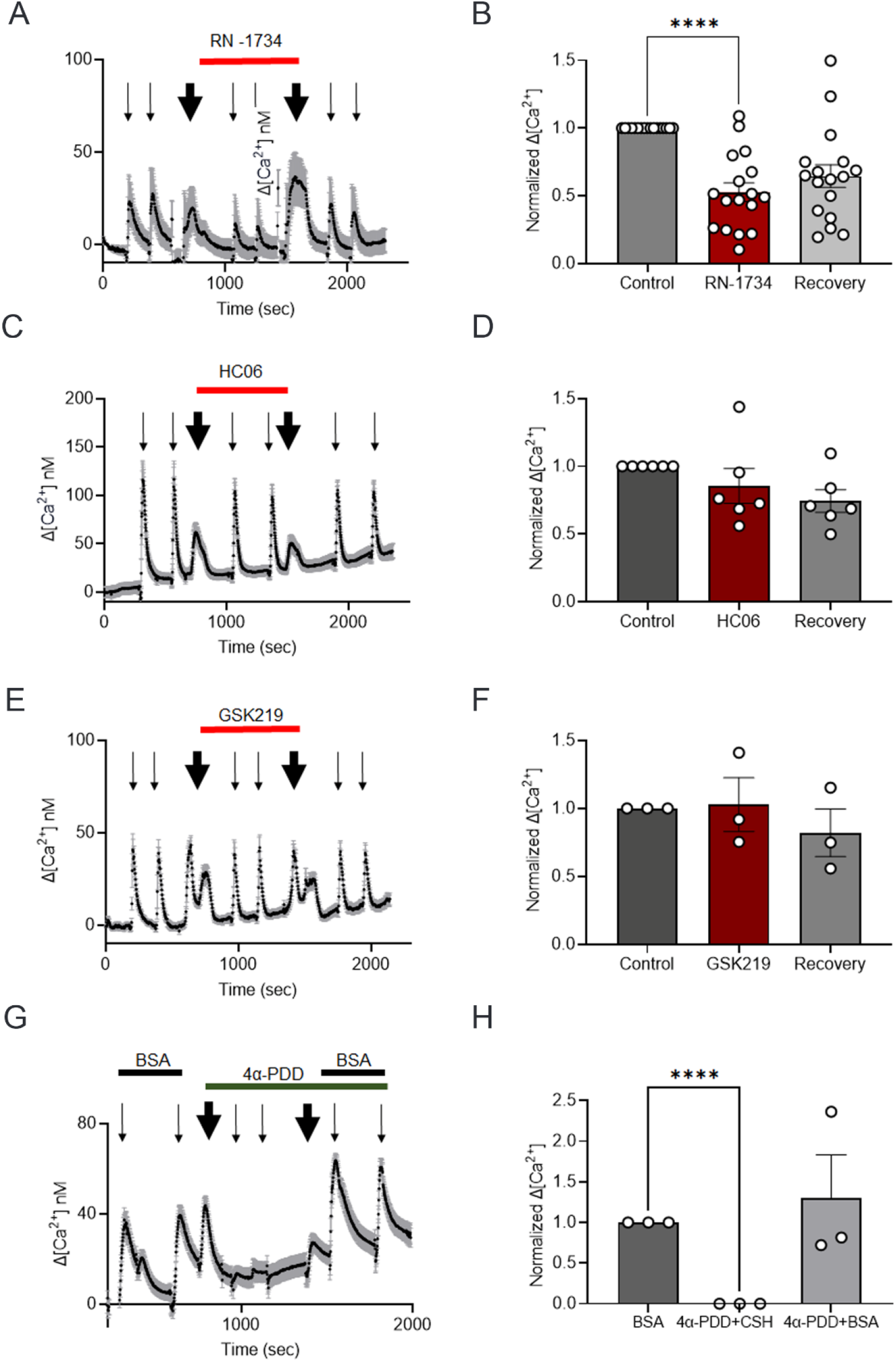
Effects of TRPV4 agonist and antagonists on flow responses. (A) Representative traces of change in [Ca2+]i showing the shear responses after acute exposure to RN-1734, a TRPV4 inhibitor. Shear forces at 5.8 dyn/cm2 for 10 seconds were applied at 300 sec intervals (thin arrows). RN-1734 was applied and removed at a flow of 0.2 ml/min for 3 minutes (thick arrows). (B) Summary of 17 independent experiments showing the partial reduction of shear-induced response by 10-30 µM RN-1734 (red bar) and the partial recovery (gray bar) upon washout. One-way ANOVA followed by Tukey’s post hoc test: ****p ≤ 0.0001. (C) Representative traces of change in [Ca2+]i showing the lack of change in shear response after acute exposure to TRPV4 antagonist HC064074. Shear forces at 5.8 dyn/cm2 for 10 seconds were applied at 300 sec intervals (thin arrows). HC064074 was applied and removed at a flow of 0.2 ml/min for 3 minutes (thick arrows). (D) Summary of 6 independent experiments showing the lack of effect on shear-induced response by 10 µM HC064074 (red bar) and the recovery (gray bar) upon washout. (E) Representative traces of change in [Ca2+]i showing the lack of change in shear response after acute exposure to 2 µM GSK2193874, another TRPV4 inhibitor. Shear forces at 5.8 dyn/cm2 for 10 seconds were applied at 300 sec intervals (thin arrows). GSK219 was applied and removed at a flow of 0.2 ml/min for 3 minutes (thick arrows). (F) Summary of 3 independent experiments showing the shear-induced response by GSK219 (red bar) and the recovery (gray bar) upon washout. (G) Representative traces of change in [Ca2+]i showing that acute exposure to TRPV4 agonist 4α-PDD in the absence of BSA did not evoke response, which was recovered upon addition of BSA. Shear forces at 5.8 dyn/cm2 for 10 seconds were applied at 300 sec intervals (thin arrows). 4α-PDD and BSA were exchanged at a flow of 0.2 ml/min for 3 minutes (thick arrows). (H) Summary of 3 independent experiments showing the lack of response in the absence of BSA despite the presence of 4α-PDD, and the recovery of response in the presence of BSA and 4α-PDD (gray bar). One-way ANOVA followed by Tukey’s post hoc test: ****p ≤ 0.0001. All data shown as mean ± SEM.

**Supplemental Figure 5:**
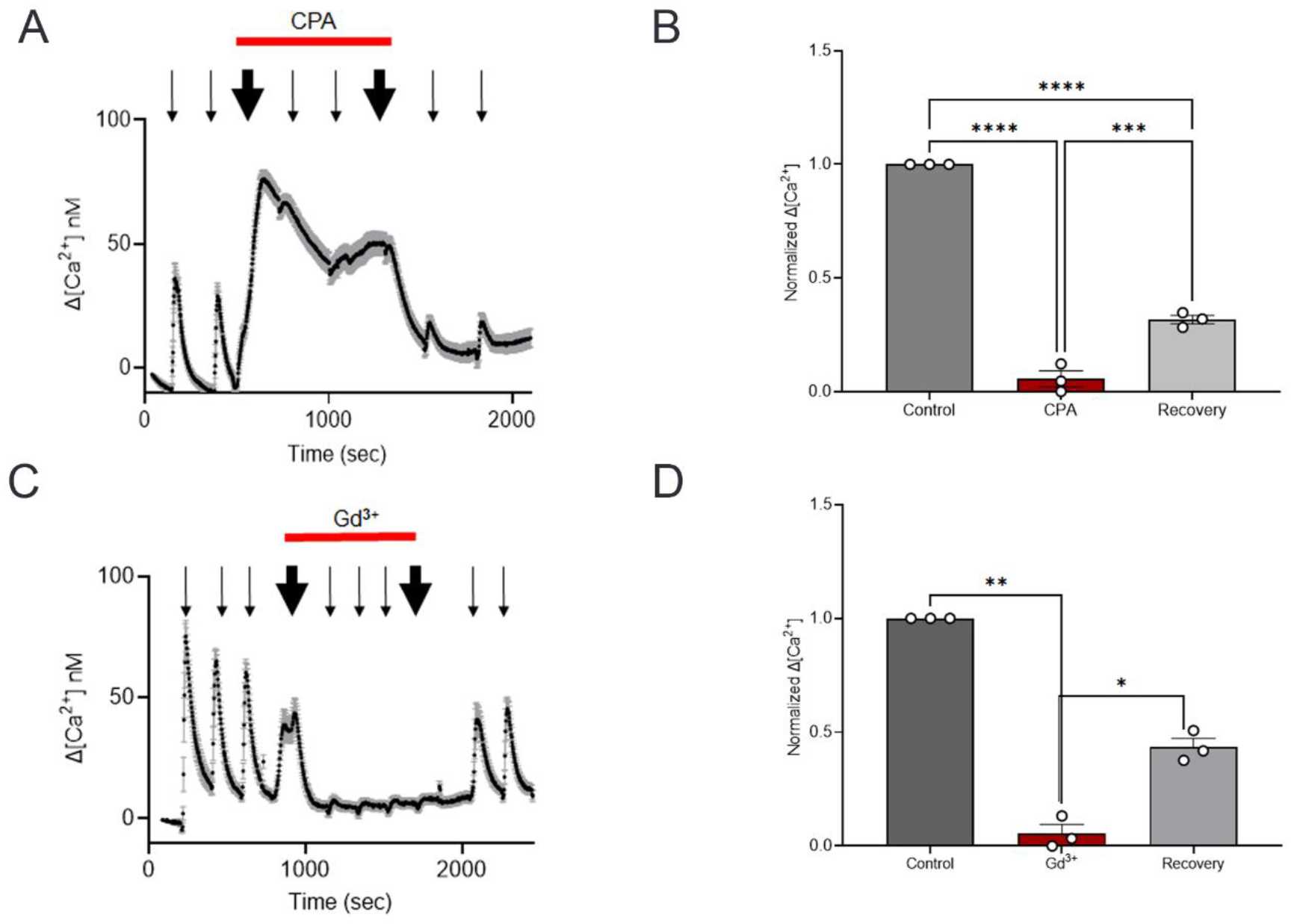
CPA and gadolinium potently inhibit calcium responses. (A) Representative traces of changes in [Ca2+]i showing the blockade of shear response after acute exposure to CPA. Shear forces at 5.8 dyn/cm2 for 10 seconds were applied at 300 sec intervals (thin arrows). CPA was applied and removed at a flow of 0.2 ml/min for 3 minutes (thick arrows). (B) Summary of 3 independent experiments showing significant reduction of shear-induced response by 10 µM CPA (red bar) and the partial recovery (gray bar) upon washout. One-way ANOVA followed by Tukey’s post hoc test: ***p<0.001; ****p<0.0001. (C) Representative traces of changes in [Ca2+]i showing the complete inhibition of shear response after acute application of gadolinium. Shear forces at 5.8 dyn/cm2 for 10 seconds were applied at 300 sec intervals (thin arrows). Gadolinium was applied and removed at a flow of 0.2 ml/min for 3 minutes (thick arrows). (D) Summary of 3 independent experiments showing significant reduction of shear-induced response by 1 mM gadolinium (red bar) and the partial recovery (gray bar) upon washout. One-way ANOVA followed by Tukey’s post hoc test: *p<0.05; **p<0.01. All data shown as mean ± SEM.

**Supplemental Figure 6:**
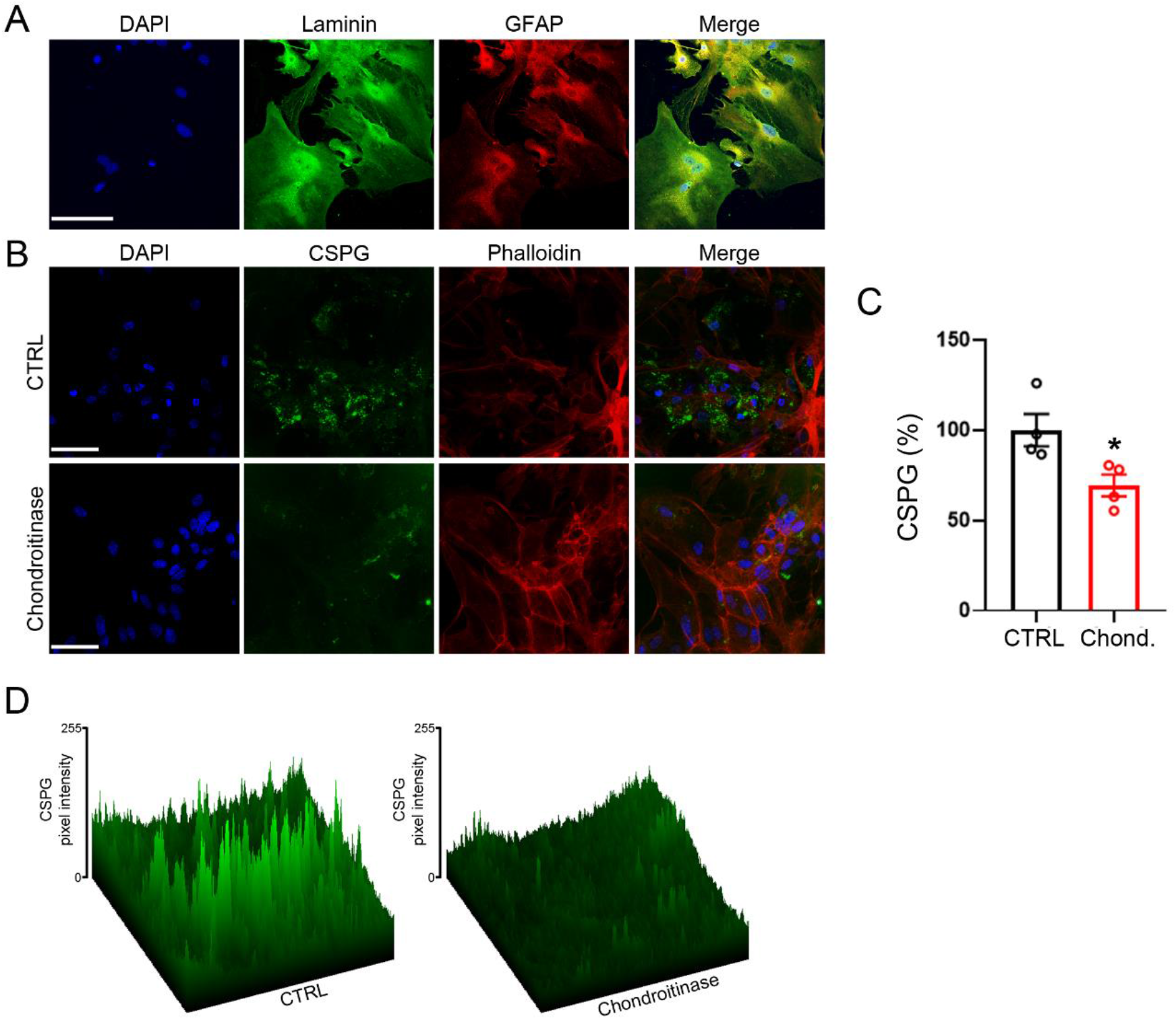
Chondroitinase ABC degrades CSPG from the cell surface of mouse astrocytes. (A) Representative single-slice confocal images of laminin (green), GFAP (red) and their merge in astrocytes. The nuclei were stained blue with DAPI. Scale bar, 50 μm. (B) Representative single-slice confocal images of CSPG (green), F-actin cytoskeleton visualized with phalloidin (red) and their merge in astrocytes. Cells were cultured in DMEM without (CTRL) and with chondroitinase ABC (Chondroitinase) for 2 h. The nuclei were stained blue with DAPI. Scale bar, 50 μm. (C) Histogram showing CSPG fluorescence intensity quantification expressed as a percentage relative to that of the control (CTRL, black), considered as 100% after 2 h treatment with chondroitinase ABC (Chond., red). Data are expressed as mean ± SEM. Four images per experiment, from a total of four experiments, were taken for each condition. Unpaired Student’s t test: *p < 0.05. (D) Reconstructed 3D surface plot images of (B). Color range from black to green indicates the relative level of fluorescence pixel intensity.

